# A formalism for sequential estimation of neural membrane time constant and input–output curve towards selective and closed-loop transcranial magnetic stimulation^⋆^

**DOI:** 10.1101/2022.04.05.487065

**Authors:** S.M.Mahdi Alavi, Fidel Vila-Rodriguez, Adam Mahdi, Stefan M. Goetz

## Abstract

**Objective:** To obtain a formalism for real-time concurrent sequential estimation of neural membrane time constant and input–output (IO) curve with transcranial magnetic stimulation (TMS).

**Approach:** First, the neural membrane response and depolarization factor, which leads to motor evoked potentials (MEPs) with TMS are analytically computed and discussed. Then, an integrated model is developed which combines the neural membrane time constant and input–output curve. Identifiability of the proposed integrated model is discussed. A condition is derived, which assures estimation of the proposed integrated model. Finally, sequential parameter estimation (SPE) of the neural membrane time constant and IO curve is described through closed-loop optimal sampling and open-loop uniform sampling TMS. Without loss of generality, this paper focuses on a specific case of commercialized TMS pulse shapes. The proposed formalism and SPE method are directly applicable to other pulse shapes.

**Main results:** The results confirm satisfactory estimation of the membrane time constant and IO curve parameters. By defining a stopping rule based on five times consecutive convergence of the estimation parameters with a tolerances of 0.01, the membrane time constant and IO curve parameters are estimated with 82 TMS pulses with absolute relative estimation errors (AREs) of less than 4% with the optimal sampling SPE method. At this point, the uniform sampling SPE method leads to AREs up to 16%. The uniform sampling method does not satisfy the stopping rule due to the large estimation variations.

**Significance:** This paper provides a tool for real-time closed-loop SPE of the neural time constant and IO curve, which can contribute novel insights in TMS studies. SPE of the membrane time constant enables selective stimulation, which can be used for advanced brain research, precision medicine and personalized medicine.

## 1. Introduction

Brain stimulation techniques, such as transcranial magnetic stimulation (TMS) or electrical stimulation (ES), can evoke neural signals through strong pulses and practically write signals into neurons in the brain, which the brain with its circuits can process similar to endogenous signals [1, 2, 3]. Experimental brain research, precision medicine, and personalized medicine desire a high level of stimulation selectivity to target only specific groups of neurons, [7, 8, 9, 10, 11]. Whereas previously stimulation selectivity was mostly achieved through improving coil focality, limits have been reached [4] and temporal aspects of pulses to exploit the neuronal activation dynamics for a further improvement of stimulation selectivity have gained attention [5, 6].

The activation dynamics are nonlinear and vary between neurons. However, typically the neural activation dynamics have a strong low-pass component, which can to some extent be approximated by a linear filter with a certain low-pass time constant. The neural membrane time constant, defined as the amount of time it takes for the linearized membrane depolarization to reach 63.2% of its final value in response to a step excitation [12], is one of the properties that significantly changes in various neurodegenerative disorders and therapeutic interventions. Its value depends on the axonal microanatomy, such as membrane resistance and capacitance, as well as the dynamics of the expressed ion channels and their sub-units [10, 13, 14].

Beyond basic neuroscientific research, identification of the neural time constant has a substantial medical value. Neurophysiological studies confirm that the membrane time constant decreases with hyperpolarization and increases with depolarization, [15]. Thus, significant attempts have been made to develop time-constant-based biomarkers for the characterization of the excitability of central and peripheral nerve systems. For instance, studies demonstrated that the time constant increases with demyelination and is significantly longer in patients with amyotrophic lateral sclerosis (ALS) and severe spinal muscular atrophy, compared to healthy controls [16, 17, 18]. After the intravenous immunoglobulin (IVIg) therapy, significant reduction of the time constant is observed in chronic inflammatory demyelinating polyneuropathy (CIDP) [19]. Sensory fibers represent significantly longer time constant than motor fibers in healthy controls and patients with alcoholic polyneuropathy, [20].

For the time constant measurement, the neural populations which result in readily detectable responses, are typically targeted, such as the primary motor cortex [21]. In the primary motor cortex, strong stimulation pulses lead to visible muscle contraction and motor evoked potentials (MEPs), which are detectable through electromyography (EMG). With recent advances in EMG instrumentation and data analytic techniques, weaker MEPs even below the noise floor of the background activity are also detectable, [22, 23, 24, 25].

Conventionally, the neural membrane time constant is measured by fitting the Lapicque or Weiss models to the S–D curve [27]. Thus, it is referred to as the S–D time constant in some papers [15, 28, 29]. The time constant estimation by using the S–D curve is performed offline, which requires stimulation at various pulse widths and computation of the motor thresholds or a similarly stable level for each pulse duration. Thus, a method which could estimate the time constant in real-time is demanding.

In [28], a model was developed based on the depolarization factor, which combines the neural membrane time constant and input–output (IO) curve in TMS. However, the depolarization factor was not computed analytically.

In this paper, the proposed integrated model is completed and discussed by analytical computation of the depolarization factor. Without loss of generality, the exact model of the controllable TMS (cTMS) pulses [11] with adjustable amplitude and width is addressed. Then, real-time sequential parameter estimation (SPE) of the neural membrane time constant and IO curve parameters is developed for both open-loop and closed-loop EMG-guided TMS.

Closed-loop TMS refers to the automatic and real-time adjustment of TMS parameters to maximize the desired plastic effects by using the brain/neural data in a feedback system [30, 31]. Closed-loop TMS is an area of active research using both closed-loop EEG-guided TMS [31, 32, 33, 34, 35] and closed-loop electromyography-guided (EMG-guided) TMS [36, 37].

In [36] and [37], two SPE methods have been developed for optimal IO curve and slope curve estimation in EMG-guided TMS, by using the Fisher information matrix (FIM) maximization. In the FIM SPE method, TMS pulse amplitudes are chosen automatically and sequentially based on prior stimulation-response pairs, in order to maximize the extracted information about the IO parameters. Thus, the FIM SPE is a closed-loop tuning method, which results in more accurate estimation of the IO curve and parameters with fewer numbers of TMS pulses, reducing the duration and discomfort of the procedure, compared to the uniform sampling method. In the uniform sampling IO curve estimation, which is an open-loop tuning method, the TMS stimuli are uniformly distributed between the minimum and maximum values, [38, 39, 40]. Despite simplicity, the uniform sampling method suffers from other drawbacks. For instance, it does not provide any information about the number of TMS pulses and their repetition. It is also not optimal in the sense that many TMS pulses might be administered that do not necessarily improve the IO curve estimation performance.

This paper presents an efficient algorithm for real-time concurrent SPE of the neural membrane time constant and IO curve parameters by using closed-loop optimal sampling and open-loop uniform sampling methods. The identifiability of the proposed integrated model, which combines the membrane time constant and IO curve, is discussed. Finally, the effectiveness of the proposed SPE methods is evaluated and discussed via extensive simulations.

### 1.1. The contributions of the paper

Compared to the previous works [11, 28], the novelties of this paper are as follows:

- Analytical calculation of the neural membrane response to cTMS pulses.
- Developing an analytical integrated model, which combines the neural activation dynamics (membrane time constant) and IO curve.
- Identifiability analysis of the proposed integrated model.
- Developing a SPE method for real-time concurrent estimation of the neural membrane time constant and IO curve parameters with open-loop uniform sampling and closed-loop optimal sampling TMS.

### 1.2. The structure of the paper

This paper is organized as follows. Section 2 develops an integrated model of the neural membrane time constant and IO curve. Section 3 first states the estimation problem. Then, the SPE method by using the optimal and uniform sampling TMS is described. Section 4 discusses the identifiability of the model parameters. Section 5 presents simulation method and results, followed by further discussions in Section 6.

## 2. The integrated model

This section develops an integrated model of the neural membrane time constant and input–output (IO) curve.

### 2.1. The TMS pulse model

In all TMS devices, an electric pulse passes through the coil, which generates a time-varying magnetic field [41]. Without loss of generality, consider the specific electric pulse waveform w(t) in a cTMS as shown in Fig. 1, which is a piece-wise signal given by [11, 42]

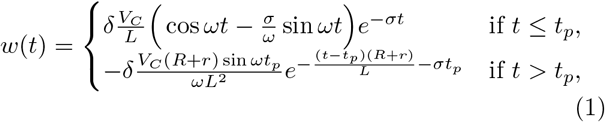

where, *V*_*C*_ denotes the normalized pulse amplitude, based on the voltage of the capacitive energy storage *C*. It is assumed that *V*_*C*_ is adjustable between *V*_*C*_(min) = 0 and *V*_*C*_(max) = 1. *t*_*p*_ is the pulse width, which is assumed to be adjustable between *t*_*p*_(min) = 10 μs and *t*_*p*_(max) = 200 μs. Other parameters include the stimulating coil *L*, the energy dissipation resistor *R*, the resistor *r* which represents the combined resistance of the capacitor, inductor and semi-conductor switch. The parameters *ω* and *σ* are defined as

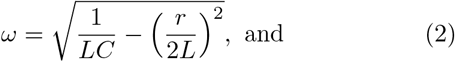

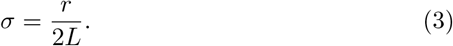

The parameter *δ* is a proportionality coefficient which describes the coupling and depends on the number of turns as well as the geometry of the coil, and the electric conductivity profile of the brain [43].

**Figure 1:**
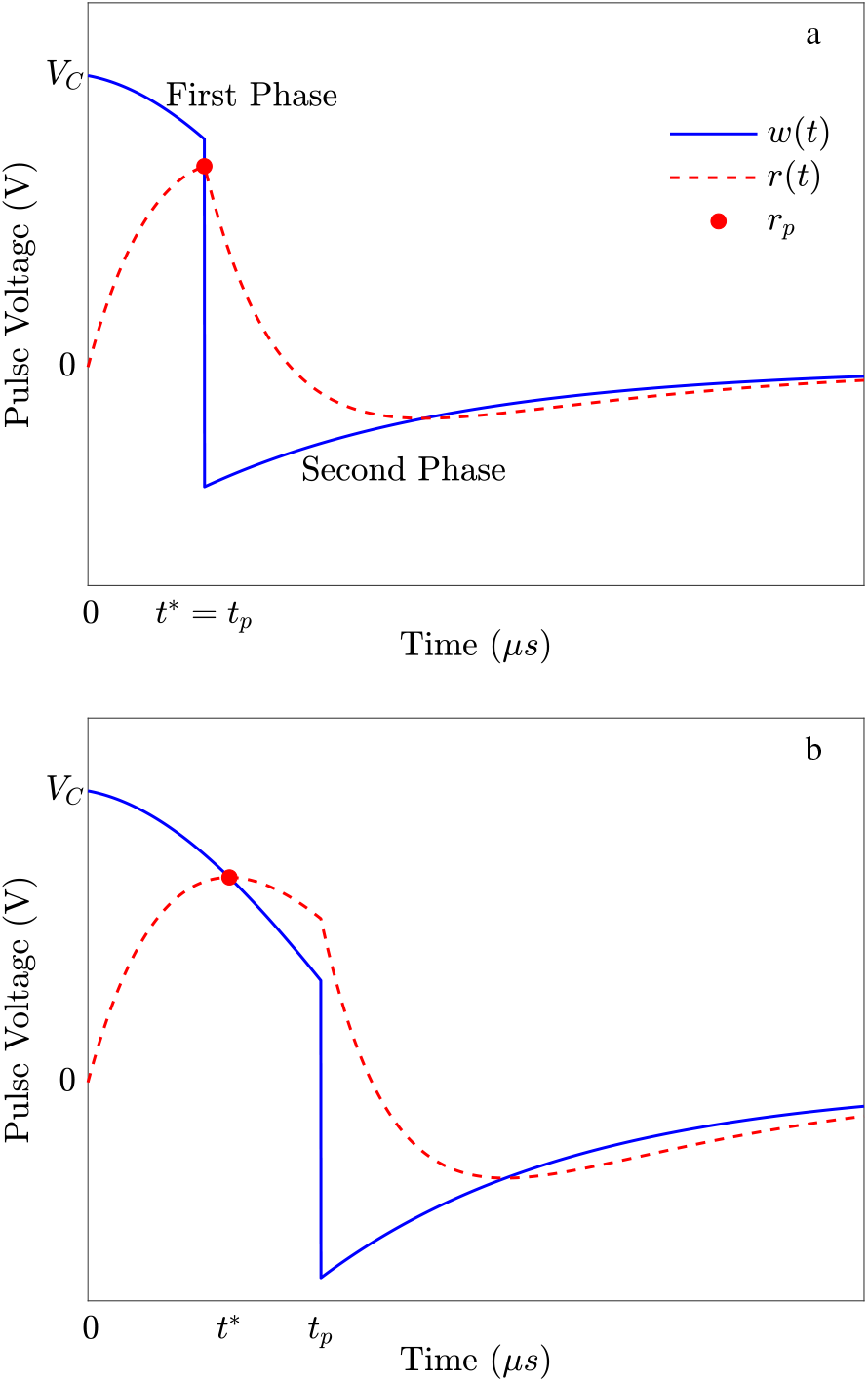
Two sample cTMS pulse waveforms with different pulse widths, and their responses of a neural membrane with the time constant *τ*. The depolarization factor *r*_*p*_ denotes the maximum (or peak) of *r*(*t*), happening at *t* = *t*^*∗*^. a) A cTMS pulse, which results in *r*_*p*_ happening at *t*^*∗*^ = *t*_*p*_. b) By increasing the pulse width, *r*_*p*_ occurs at *t*^*∗*^ < *t*_*p*_.

Following the original cTMS design and without loss of generality, the following parameter values are used in this paper [11]: *L* = 16 μH, *C* = 716 μF, *R* = 0.1 Ω, *r* = 20 mΩ, *δ* = 3.2 *×* 10^*−*6^ (V/m)(A/s).

**Remark 1**. *In conventional TMS devices, the pulse width is fixed [44, 41]. Thus, the proposed modeling is also applicable to them as a special case just as it is to more advanced TMS technology with more flexible pulse control [45, 46, 47]*.

### 2.2. The neural membrane model and response

The neural membrane depolarization response *r*(*t*) is computed by the convolution of the administered pulse *w*(*t*) and the neural membrane dynamic represented by its impulse response *h*(*t*), which is approximated by a first-order low-pass filter with time constant *τ* as follows [28, 48, 44]

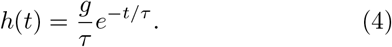

Different first-order time constants have been reported in the literature, e.g. 196 μs in [28], (152 *±* 26) μs in [48], and even more than 200 μs in [29]. This emphasizes the need for a method for the estimation of *τ*. The aim of this paper is to present a method for the sequential estimation of *τ*. The parameter *g* represents the coil-to-neuron coupling gain, which depends on factors such as the coil-to-cortex distance, head size, and neural population type and orientation relative to the induced electric field. The estimation of *g* is beyond the scope of this paper. This paper assumes that *g* is a fixed value between 30 and 50.

Given *w*(*t*) and *h*(*t*), the neural membrane depolarization response *r*(*t*) is obtained as

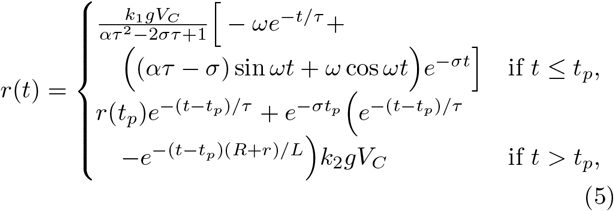

where, *r*(*t*_*p*_) is the neural membrane response at *t* = *t*_*p*_, and

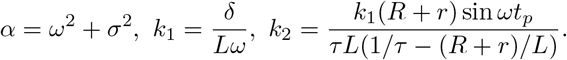

Details of the formalism are given in Appendix A.

The depolarization factor *r*_*p*_ denotes the maximum or peak point of *r*(*t*) defined as

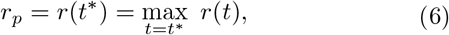

where, *t*^*∗*^ is the peak time.

Fig. 1 shows two cTMS pulse waveforms with different pulse widths, and their corresponding responses of a first-order neural membrane approximation with the time constant *τ*. It is observed that the maximum or peak point *r*_*p*_ occurs either at the end of the pulse (*t*^*∗*^ = *t*_*p*_) or before the end of the pulse (*t*^*∗*^ < *t*_*p*_).

The critical pulse width 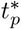 is defined as the pulse width for which *r*_*p*_ occurs at *t*^*∗*^ = *t*_*p*_ for all pulse widths shorter than that, and *r*_*p*_ occurs at *t*^*∗*^ < *t*_*p*_ for all pulse widths longer than that. The values of *t*^*∗*^ and 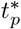 depend on the the pulse parameters and *τ*.

Fig. 2 shows the relationship between 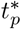 and *τ* for the cTMS pulse parameters given in Section 2.1. This figure is obtained by manually solving the maximization problem (6), for the time constants from *τ* = 10 μs to *τ* = 215 μs with the step size of 5 μs. It is seen that the critical pulse width increases exponentially with the membrane time constant. The next section discusses the importance of the critical pulse width on identifiability of the membrane time constant.

**Figure 2:**
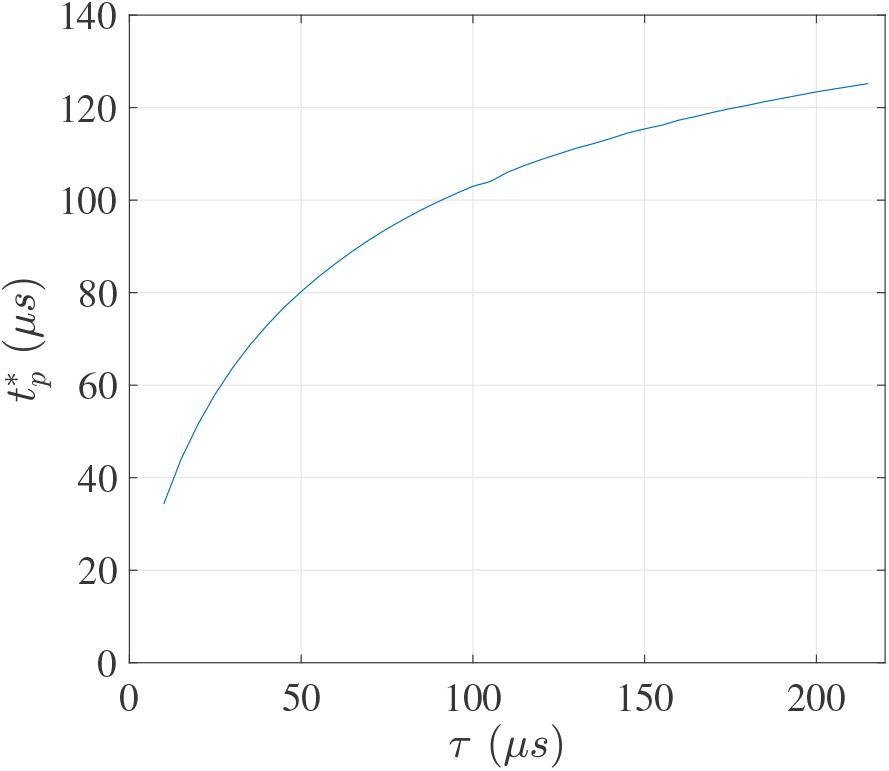
The relationship between the critical pulse width 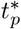 and the membrane time constant *τ*.

### 2.3. The input–output curve model

As shown in the literature, the IO curve, characterizing the relationships between the MEPs and pulse amplitudes would be sigmoidal for a fixed *t*_*p*_ when TMS pulses are administered with the same pulse-width *t*_*p*_ [28, 36, 49]. On the other hand, an MEP is generated when the depolarization factor *r*_*p*_ reaches a threshold.

Whereas earlier studies used to describe MEPs and IO curves linearly, research has demonstrated that IO curves reside in the log domain and show intricate variability characteristics [49, 50]. The variability can be normalized by log-transformation in the first approximation [51, 52]. Accordingly, the peak-to-peak MEP measurements *V*_pp_ are log-transformed through *y* = log (*V*_pp_), and the IO curve is modeled by the following sigmoid function

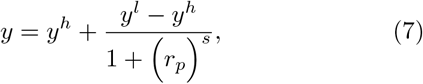

where *θ*_1_, *θ*_2_, *θ*_3_, and *θ*_4_ denote the lower plateau, upper plateaus, mid-point and slope of the IO curve, respectively.

The sigmoid model (7) is parameterized in the standard format, *y* versus *V*_*C*_, as

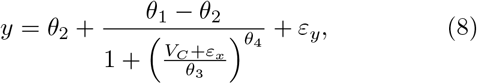

where

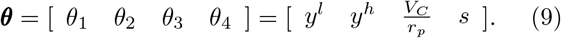

In order to reproduce the intricate statistical distributions observed in measurements, the random noises *ε*_*x*_ and *ε*_*y*_ are added along x and y axes to address the neurophysiological and electromagnetic uncertainties and variabilities, such as trial-to-trial variability, excitability fluctuations, variability in the neural and muscular pathways, positioning fluctuations, and physiological and measurement noise, [53].

It is re-called that the neural membrane time constant, *τ*, appears in *r*_*p*_ and consequently in *θ*_3_.

Figure 3 illustrates this integrated model structure for a cTMS pulse *w* to MEP *y* in a block diagram.

**Figure 3:**
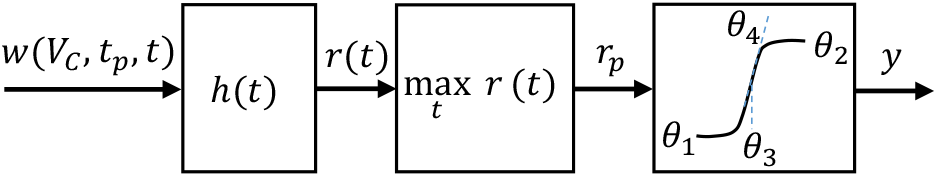
The functional block diagram of the integrated model of the neural membrane time constant and input–output curve from the cTMS pulse *w* to MEP *y*.

## 3. The Proposed Sequential Parameter Estimation Methods

### 3.1. Problem statement

As the key problem, this paper aims to concurrently estimate the neural membrane time constant *τ* and the IO curve parameters *θ*_*i*_, i = 1, …, 4 with TMS.

This section proposes a SPE method based on the closed-loop optimal sampling and open-loop uniform sampling principles.

### 3.2. Outline of overall procedure

SPE starts with sampling from the baseline, administering three initial random pulses, measuring the corresponding MEPs, and computing the initial estimate of the IO curve parameters and time constant. The next TMS pulses are then chosen based on the uniform or optimal sampling methods. After the administration of each pulse, the corresponding MEP is measured, and the estimation of the IO curve and time constant is updated. The process continues until a stopping rule is satisfied or the maximum number of pulses is reached. Thus, the SPE method consists of four stages: initialization, selection of next TMS pulse, estimation update, and checking the stopping rule.

Notably, the entire estimation processes is a fully automatic online closed-loop procedure. It comprises the computation of the next TMS pulse with its specific parameters needed to extract most information in the next step via administration of the pulse to the EMG recording of the MEP response to the pulse and data analysis.

### 3.3. Initialization

In the initialization stage, the baseline data

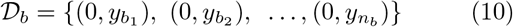

is collected. The baseline data is the EMG data in the absence of TMS pulse *V*_*C*_ = 0. In (10), *n*_*b*_ denotes the number of samples taken from the baseline. Then, three random TMS pulses with pulse strengths 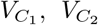, and 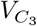 are administered and the corresponding MEPs *y*_1_, *y*_2_, and *y*_3_ are measured. An initial estimate of the IO curve and time constant parameters is obtained by fitting the model (8) to the data set of

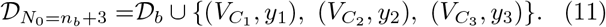

The estimates of the IO curve parameters and time constant after the administration of the n-th stimulus 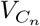 are named 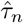 and 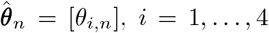. At the end of the initialization stage, the procedure obtained 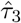 and 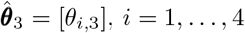.

### 3.4. Selection of the next TMS pulse

In the next stage, the next TMS pulse is chosen by uniform or optimal sampling. In uniform sampling, the TMS stimuli are uniformly distributed between *V*_*C*_(min) and *V*_*C*_(max) and could be administered randomly or in an order, e.g., from weak to strong stimuli. However, studies show hysteresis due to strong correlation between subsequent responses, when pulses are administrated in an order [54, 55].

In the optimal sampling method, 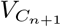 is determined by solving the optimization problem

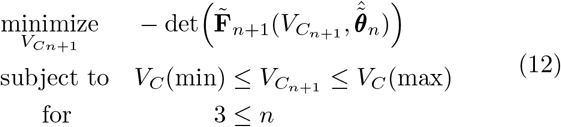

where,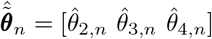, 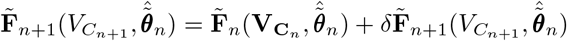, and 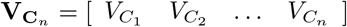, following a fundamental concept suggested previously [36]. The matrices 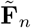 and 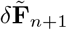 are given in Appendix B.

### 3.5. Estimation update

After each pulse 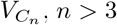, the corresponding MEP *y*_*n*_ is measured, and the data set is updated per

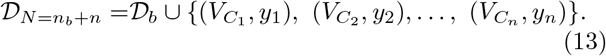

The estimation of 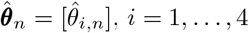 is then updated by fitting the curve of (8) to the data set of 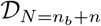. In order to mitigate the otherwise skewed variability, the curve fitting is performed on the logarithmic scale [29, 28]. After obtaining 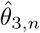, the estimation of 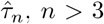 is obtained by solving

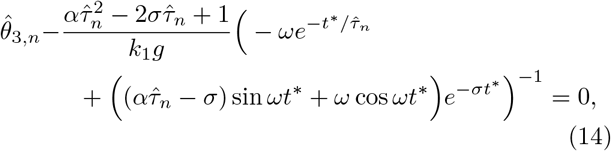

where, *t*^*∗*^ is the time at which the linearized neural membrane depolarization reaches its maximum as described in Section 2.2.

A previously developed bad-fit detection and removal technique is employed as described in the literature to improve curve fitting and solving the nonlinear equation (14) [36, 37].

### 3.6. Termination of checking the stopping rule

The SPE is terminated when the following convergence criteria are satisfied for *T* successive times

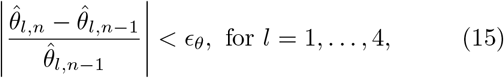

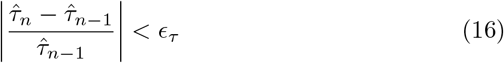

where, *ϵ*_*θ*_ and *ϵ*_*τ*_ denote the estimation tolerance of ***θ*** and *τ*, respectively. To reduce premature terminations, *T* > 1 is strongly suggested as discussed in the literature, [36].

If the aforementioned stopping rule is not satisfied, the process will continue until the maximum number of pulses is administered.

## 4. Identifiability Analysis

After administering the n-th cTMS pulse 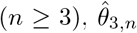 is obtained through curve regression. In order to continue the SPE method, the nonlinear equation (14) must be solvable. Thus, the identifiability of the IO curve and membrane time constant depends on the solution of (14).

If *t*^*∗*^ is known, 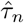 will be the only unknown parameter in (14), which can be found by using numerical methods. Based on the discussions in Section 2.2, *t*^*∗*^ is known only if *t*_*p*_ is less than the critical pulse width 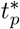. Thus, the short pulse width is recommended from the identifiability perspective. In this case, *r*_*p*_ occurs at the end of the pulse width, *t*^*∗*^ = *t*_*p*_, the nonlinear equation (14) is solvable, and the parameters 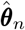 and 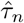 are identifiable.

Remaining is the solution of the nonlinear equation (14) with the known *t*^*∗*^ and unknown 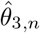. Fig. 4 displays 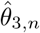 versus 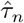 for different *t*^*∗*^s. For *t*^*∗*^ = 30 *μs*, a unique 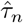 is obtained for almost all 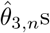. For *t*^*∗*^ = 60 *μs*, two 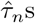 are obtained for 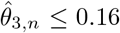, and there will be a unique solution for 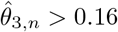. By increasing the *t*^*∗*^ value, the region that results in two 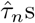 increases. The acceptable time constant is obtained based on the neurophysiological or microanatomical characteristics of the neural membrane.

**Figure 4:**
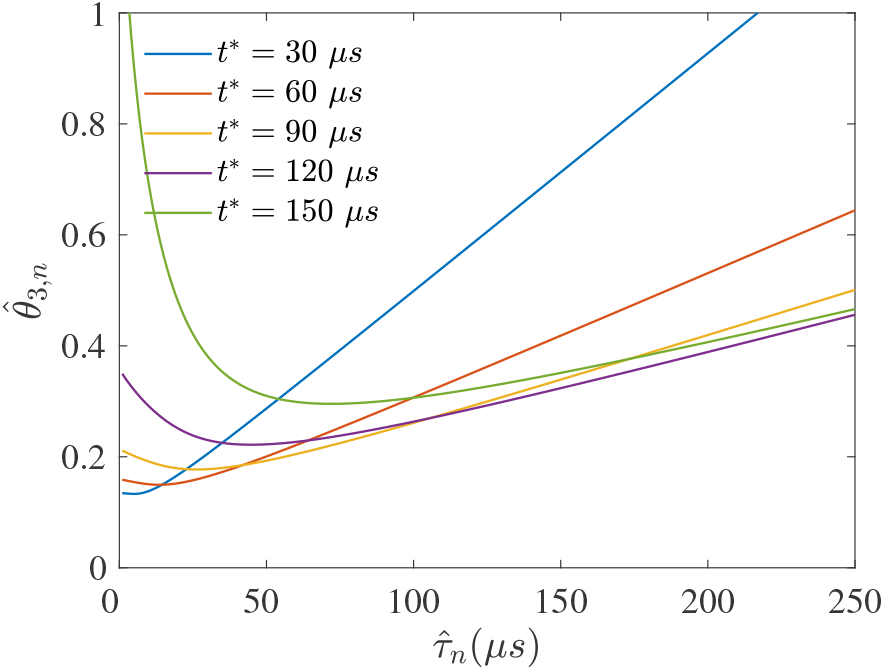
Identifiability of 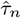.

## 5. Simulation Method and Results

### 5.1. Simulation method

Consider 10,000 stimulus–response pairs, shown by ‘*×*’, in Fig.5-a, generated by using the integrated model (8), with the following randomly chosen neural time constant and IO curve parameters [53]

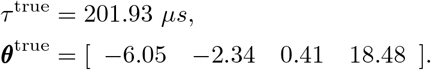

From Fig. 2, the critical pulse width for *τ* ^true^ = 201.93 is 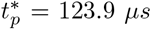. Thus, the integrated model is identifiable for pulse widths less than 123.9 *μs*. The pulse width is randomly fixed at *t*_*p*_ = 100 *μs*. 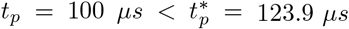, thus, *t*^*∗*^ = *t*_*p*_ = 100 *μs*. The gain g is arbitrarily set to 40.

**Figure 5:**
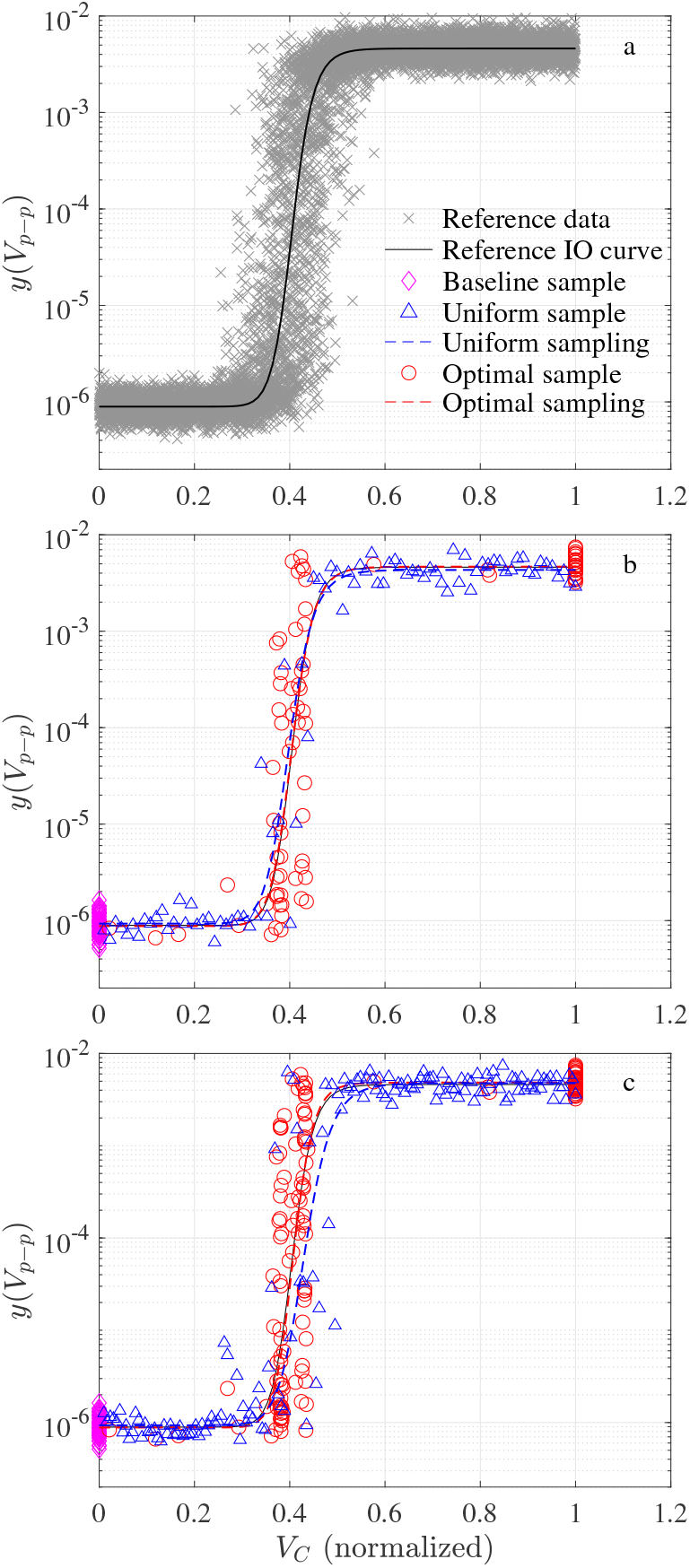
Sample simulation run: (a) reference data and IO curve, and estimation by using the optimal (FIM SPE) and uniform sampling methods at *n* = *n*_f_ = 82 (b) and at *n* = *n*_max_ = 150 (c).

The random noises *ε*_*x*_ and *ε*_*y*_ are generated by zero-mean normal functions with standard deviation of 0.05 and 0.1, respectively. The *y* axis denotes the MEP’s peak-to-peak value used in the simulation. The solid line in Fig. 5-a shows the true IO curve in the absence of *x* and *y* variabilities.

The problem in this section is to estimate the neural time constant and IO curve parameters by using the proposed optimal and uniform sampling SPE methods. The stopping rule is set up with *T* = 5, i.e., five times consecutive satisfaction of the convergence criteria in (15) and (16) with *ϵ*_*θ*_ = *ϵ*_*τ*_ = 0.01. The maximum number of TMS pulses is set to *N*_max_ = 150. Fifty samples are taken from the baseline for both methods.

The simulation is run in Matlab. The estimation update runs a trust-region regression algorithm with lower and upper limits on the parameter vector per

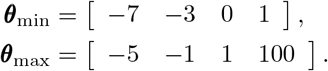

The FIM optimization of Eq. (12) and the estimation of Eq. (14) are solved by using ‘fmincon’ with ‘GlobalSearch’-‘interior-point’ algorithms.

### 5.2. Simulation results

The optimal sampling SPE method satisfied the stopping rule at *n* = *n*_*f*_ = 82, while the uniform sampling SPE did not fulfill the stopping rule. Figures 5-b and c show the acquired samples and estimated IO curves by the optimal and uniform sampling SPE methods at *n*_*f*_ = 82 and *N*_*max*_ = 150, respectively. As discussed in [36], it is seen that the optimal sampling SPE method administers stimuli, mainly from three regions, which contain the maximum information of the IO curve. These regions include two sectors from the slope area and one sector at around *x*_max_ = 1. In contrast, the uniform sampling administers TMS stimuli across the entire range of *x* = 0 to *x* = 1. Figure 6 displays the estimates of *θ*_*i*_, *i* = 1, …, 4 with increasing pulse number. Figure 7, in turn, provides the corresponding *τ*.

**Figure 6:**
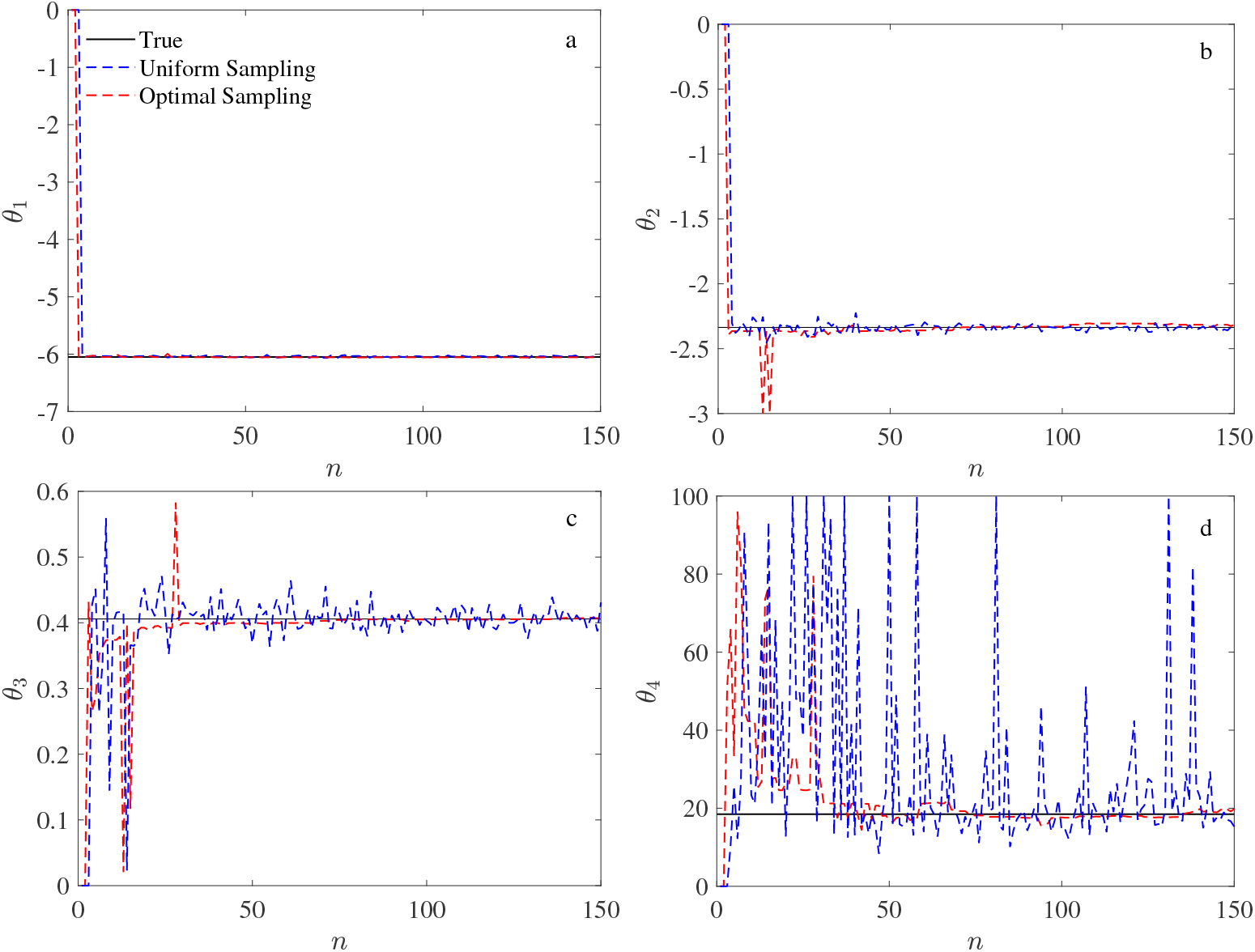
Estimated IO curve parameters: (a) lower plateau, (b) upper plateau, (c) mid-point, and (d) slope.

**Figure 7:**
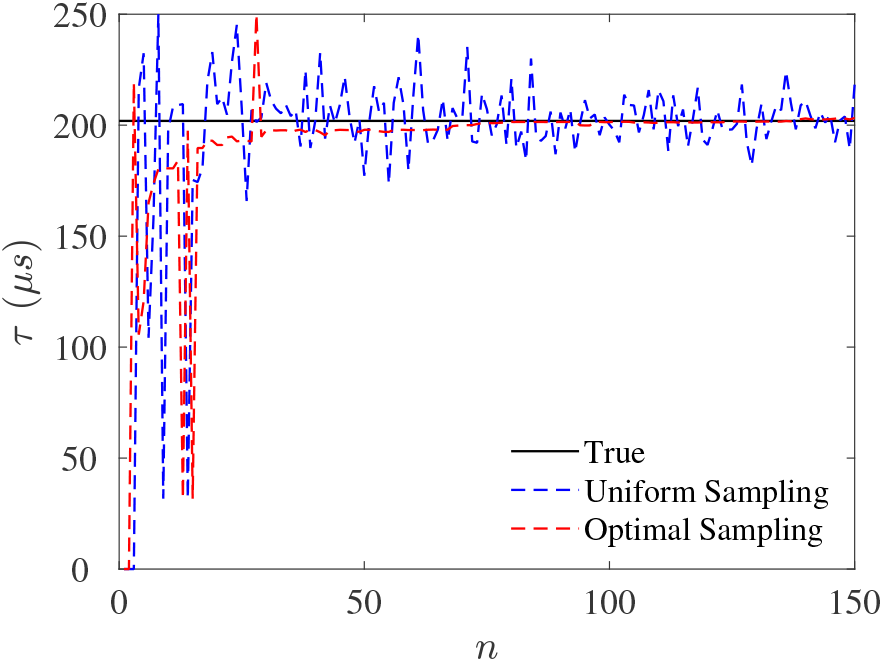
Estimated neural time constant.

When the optimal sampling SEP satisfies the stopping rule at *n* = *n*_*f*_ = 82, *θ*_1_, *θ*_2_, *θ*_3_, *θ*_4_, and *τ* are estimated with the absolute relative estimation errors of 0.13%, 0.24%, 0.18%, 3.74%, and 0.24%, respectively. In contrast, the uniform sampling method results in AREs of 0.29%, 1.22%, 1.78%, 15.62%, and 0.23% at *n* = *n*_*f*_ = 82 for *θ*_1_, *θ*_2_, *θ*_3_, *θ*_4_, and *τ*, respectively. It is seen in Figures 6 and 7 that the uniform sampling method leads to larger estimation variations.

## 6. Discussion

The integrated model (8) contains the information of the membrane activation dynamics, TMS pulse parameters, and IO curve parameters. It also considers the neurophysiological and technical uncertainties and variabilities which exist along x and y axes. Therefore, the integrated model (8) is more comprehensive than the models employed in [28, 36, 37, 38, 39, 40]. The depolarization factor or the maximum (peak) of the depolarization response occurs either at the end or before the end of the TMS pulse. For solving Eq. (14), the pulse duration should be chosen such that the depolarization factor happens at the end of the pulse or equivalently, the pulse duration is less than the critical pulse width. If the range of the time constant is approximately known, for instance from the literature, the approximate critical pulse width can be obtained from Fig. 2. If there is no prior information about the time constant, short pulse duration is recommended.

This paper assumes that the coupling gain g is fixed. The estimation of the coupling gain and time constant requires concurrent estimation of more interleaved IO curves, which is beyond the topic of this paper.

The proposed optimal and uniform sampling based techniques for SPE of the membrane time constant are run in real-time, which is an advantage compared to the conventional methods based on the strength-duration curve, where the motor threshold needs to be calculated offline at several pulse widths. Also, the proposed techniques concurrently estimate the IO curve parameters.

It should be noted that previous comparisons in [36], clarifying the differences between the uniform and optimal sampling SPE methods are still valid here. With reference to the results of 10 177 simulation runs in [36], the low and high plateaus and mid-point (*θ*_1_, *θ*_2_, and *θ*_3_, respectively) are estimated with a median relative error of less than 2% for *T* = 5. The slope *θ*_4_ is estimated with the median relative error of less than 20% for *T* = 5. Note that the neural time constant *τ* relates only to *θ*_3_ through (9). Thus, the estimation error of *τ* is proportional to the estimation error of *θ*_3_.

## 7. Conclusions and future works

This paper describes a novel formalism for sequential parameter estimation (SPE) of the neural time constant through the input–output (IO) curve and its parameters in transcranial magnetic stimulation (TMS). Analytical calculation of the depolarization factor was discussed. An integrated model was developed based on the depolarization factor, relating the neural recruitment curve and time constant. The identifiability of the proposed integrated model was analyzed. The formalism was linked with uniform and optimal (FIM-based) sampling principles. The results demonstrate estimation of the neural time constant through the IO curve parameters in the presence of x and y variabilities. Without loss of generality, this paper focused on a specific case of cTMS pulses. The method is directly applicable to other cTMS pulse shapes with no new point of principals. The development of the proposed SPE method to higher-order and/or nonlinear membrane dynamical models appears as an auspicious mathematical challenge for further work though it will require substantially more data points and does not have any medical support yet. Practical implementation of the proposed SPE method remains for the future work.

## 8. Appendix A

The state-space representation of the neural membrane dynamic *h*(*t*) is given by

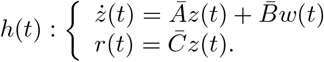

where *z* and *r* are the membrane state and response with

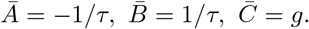

Given the system matrices 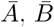, and 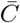, the membrane response *r*(*t*) is computed by using

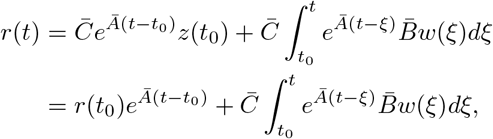

where *t*_0_ is the initial time [56].

Within the first phase of *w*(*t*), i.e., *t ≤ t*_*p*_, *t*_0_ = 0, *r*(0) = 0, the membrane response is computed as

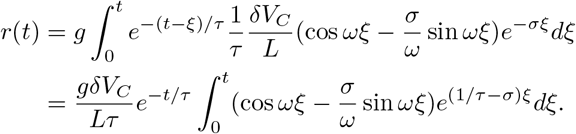

By defining *a* = 1/*τ − σ*, the integration by parts results in

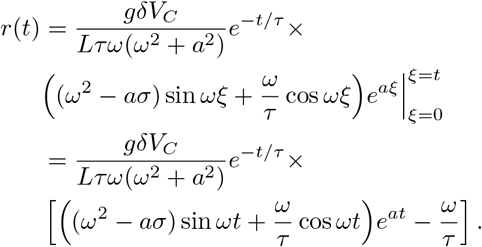

Defining *k*_1_ = *δ*/(*Lω*) and *α* = *ω*^2^ + *σ*^2^ results in the part of Eq. (5) for *t ≤ t*_*p*_.

Within the second phase of *w*(*t*), i.e., 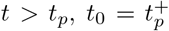, where 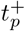 denotes a time slightly after *t*_*p*_. By using the approximation 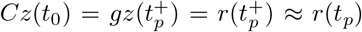, the membrane response is computed as follows,

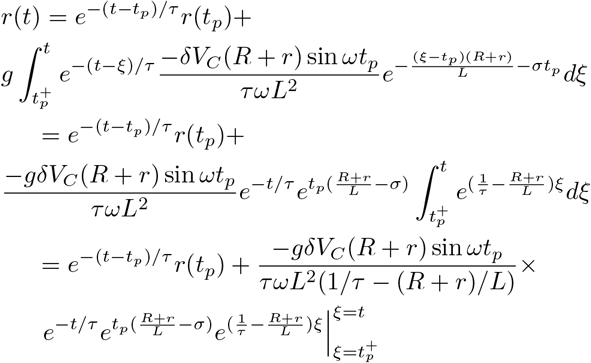

With the approximation 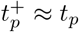,

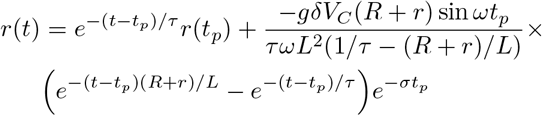

Defining 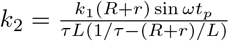 leads to the part of Eq. (5) for *t* > *t*_*p*_.

## 9. Appendix B

The matrices in the FIM optimization problem per Eq. (12) are computed as follows. The matrix 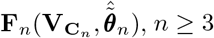 is given by

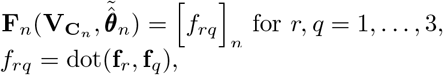

where

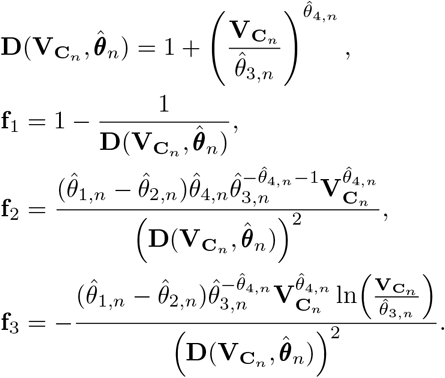

The bilinear form dot(**f**_*r*_, **f**_*q*_) returns the scalar product of the vectors **f**_*r*_ and **f**_*q*_.

The matrix 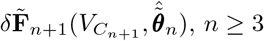 is computed per

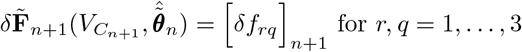

where

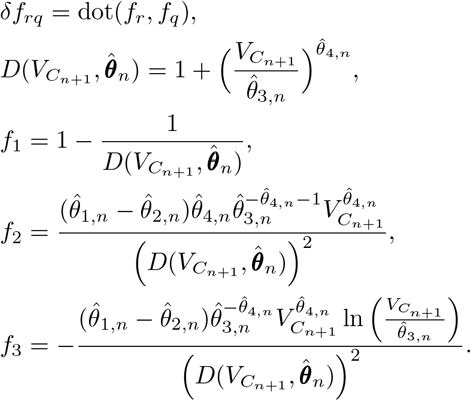

